# Perineuronal nets reflect a continuum of fast-spiking specialization in adult parvalbumin interneurons

**DOI:** 10.64898/2026.03.31.715401

**Authors:** Sverre Grødem, Guro Helén Vatne, Kristian Kinden Lensjø, Kosio Beshkov, Mads Lønø, Torkel Hafting, Marianne Fyhn

## Abstract

Perineuronal nets (PNNs) preferentially enwrap parvalbumin (PV) interneurons, regulating plasticity and circuit function. The molecular differences between PNN-positive and PNN-negative PV neurons remain unknown. We combined Xenium spatial transcriptomics with PNN-labeling in adult mouse cortex (378,349 cells) and found that 97% of PNNs enwrap PV neurons; substantially higher than previous estimates. PNN status reflected a transcriptional continuum rather than discrete subtypes. A classifier trained on Xenium data (AUC = 0.87) applied to Allen Brain scRNA-seq data, enabled genome-wide analysis of 34,326 cortical PV neurons. PNN-positive PV neurons expressed mature fast-spiking markers: Kv3 channels, mature NMDA subunits (Grin2a), fast-kinetic GABA-A receptors (Gabra1), oxidative phosphorylation genes, and gap junctions (Gjd2). PNN-negative PV neurons expressed neuropeptides and GABA-A subunits typical of Sst interneurons, suggesting a transcriptomic boundary between cell types and potentially elevated plasticity. This establishes PNN status as a molecular correlate of PV cell specialization, with implications for therapeutic strategies targeting cortical plasticity.

## INTRODUCTION

Cortical circuits rely on a precise balance of excitation and inhibition, orchestrated largely by a diverse population of GABAergic interneurons^1,2^. Among these, fast-spiking parvalbumin-expressing (PV) cells are critical for generating gamma oscillations, shaping sensory responses, memory consolidation, and network stability^3–7^. A subset of PV neurons are enwrapped by perineuronal nets (PNNs), specialized extracellular matrix structures that aggregate around the soma and proximal dendrites^8^. PNNs are composed of hyaluronan, chondroitin sulfate proteoglycans (particularly aggrecan), link proteins, and tenascin-R, forming a dense lattice that physically constrains synaptic remodeling^8,9^. While PNNs form during postnatal development and contribute to critical period closure, they persist throughout adulthood, stabilizing synaptic connections and restricting plasticity^6,10,11^. However, even in the fully mature brain, not all PV neurons bear PNNs, and the molecular distinctions between these co-existing populations remain poorly characterized. Understanding the molecular basis of this heterogeneity could reveal mechanisms underlying differential circuit plasticity and inform conditions where PNN disruption has been implicated, including schizophrenia, Alzheimer’s disease, and epilepsy^12^.

Previous histochemical studies have reported that 20–30% of cortical PNNs surround non-PV neurons, suggesting heterogeneity in PNN-enwrapped cell types^13–15^. However, Pvalb immunostaining has limited sensitivity, and can potentially miss neurons with low Pvalb protein levels. Recent transcriptomic atlases have revealed extensive heterogeneity within the adult PV cell-types, resolving multiple transcriptionally distinct subtypes^16–18^. This includes neurons co-expressing Pvalb with neuropeptides such as somatostatin (Sst). Notably, Pvalb-Sst co-expressing neurons have been reported to lack PNNs in a limited patch-seq dataset^19^. Whether PNN status more broadly maps onto transcriptomic heterogeneity within the PV population, and what molecular programs distinguish PNN-positive from PNN-negative neurons in adulthood, remains unknown. To address these questions, we combined spatial transcriptomics (Xenium, 297 genes, Table S1.1) with Wisteria Floribunda Agglutinin (WFA) labeling, which robustly labels PNNs, on the same tissue sections from adult mice (7 months old). This revealed that 97% of PNN-positive cells belong to PV subclasses, substantially higher than immunohistochemical estimates^13^. Within the cortical PV basket cell population, we found that PNN status maps as a transcriptional continuum, rather than a discrete subtype. Next, we trained a classifier on Xenium data (nested cross-validation, AUC = 0.87) that predicted PNN status from a compact gene signature. Applying this classifier to Allen Brain scRNA-seq data^16^ enabled genome-wide differential expression analysis of 34,326 PV neurons.

Genome-wide analysis afÏrmed that PNN status reflects a transcriptional continuum and revealed its functional basis: the molecular machinery underlying fast-spiking physiology and circuit integration. PNN-positive neurons display a canonical mature, fast-spiking signature: Kv3 channels essential for rapid repolarization^20^, Grin2a over developmental Grin2b^21^, GABA receptor subunits conferring faster inhibitory kinetics^22^, elevated oxidative phosphorylation machinery, and gap junction coupling expression via Gjd2^23^. PNN-negative PV neurons express neuropeptides typically associated with other interneuron classes (Sst, Tac1, Npy) and GABA receptor subunits characteristic of Sst interneurons^22,24,25^, suggesting that they occupy a transcriptional boundary between PV and Sst identities. These findings indicate that PNN status provides a molecular readout of position along a specialization axis persisting into adulthood, with implications for experience-dependent remodeling and PNN-targeted therapeutics.

## RESULTS

### Spatial transcriptomics with PNN labeling reveals near-exclusive association with PV in-terneurons

To characterize the transcriptomic identity of PNN-bearing neurons, we applied Xenium spatial transcriptomics (10x Genomics, 297-gene panel) to sagittal brain hemi-sections (1.5-2mm ML) from adult mice (P220, n = 5 animals, 2 sections per animal), followed by fluorescent WFA labeling on the same tissue to label PNNs (Figure 1A). The sections contained primarily isocortex and hippocampus, with portions of striatum and thalamus (Table S1.6). After quality filtering, we retained 378,349 cells, which were classified into cell types using the Allen Brain MapMyCells pipeline^16,26^, and brain areas by reference atlas alignment^27^. PNNs were manually detected based on WFA labeling across all slices by an annotator blinded to transcriptomic data.

**Figure 1.**
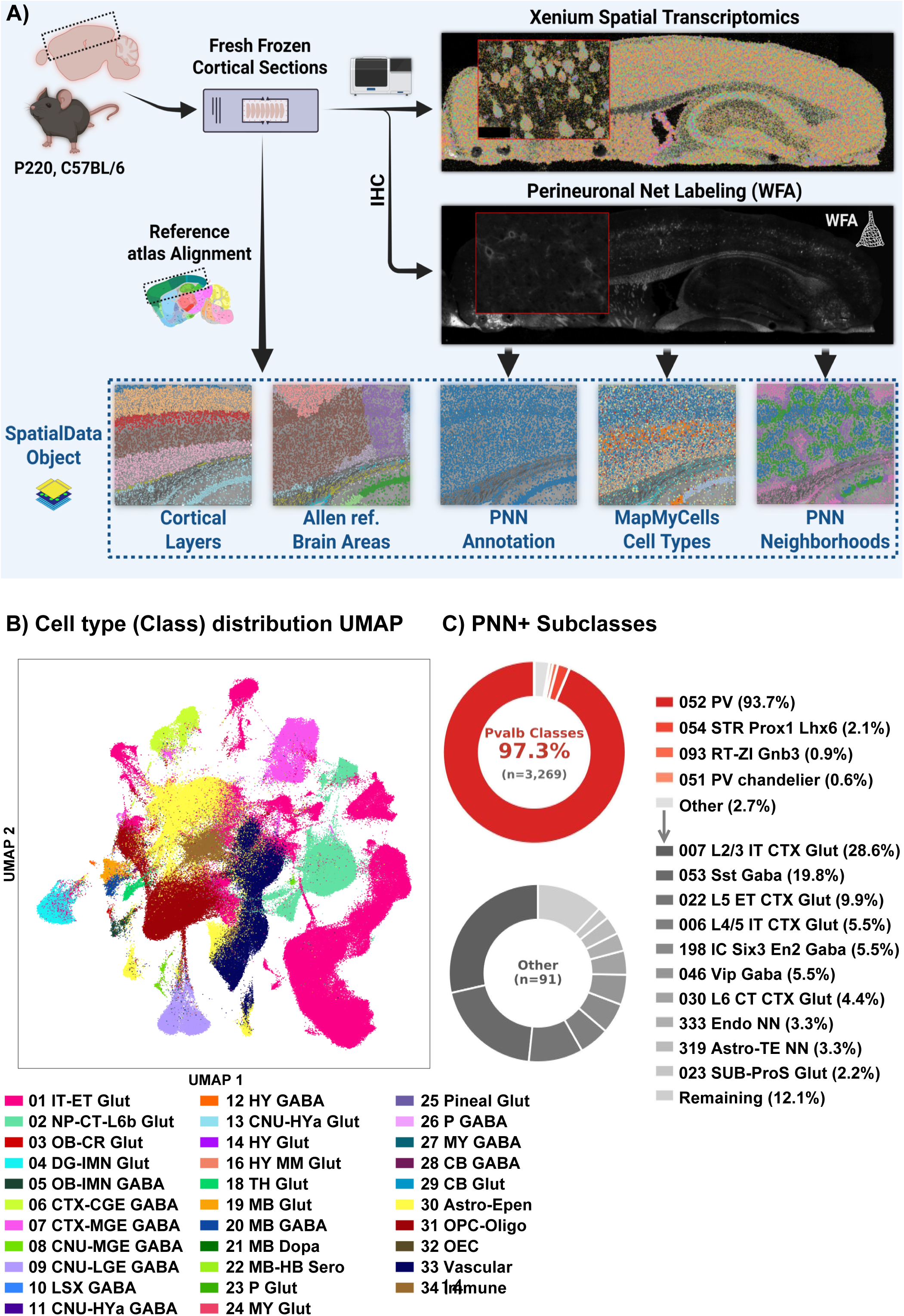
Spatial transcriptomics and perineuronal net mapping across the adult mouse cerebrum. **(A)** Overview of the experimental pipeline. Coronal brain sections from P220 mice were pro-cessed with the 10x Genomics Xenium in situ transcriptomics platform, yielding spatially resolved transcript maps. The same sections were stained with Wisteria floribunda agglutinin (WFA) to label perineuronal nets (PNNs). Xenium tissue sections were aligned to the Allen Mouse Brain Common Coordinate Framework reference atlas using the QUINT workflow (QuickNII/VisuAlign). Cell-type annotations were obtained via the Allen Brain MapMyCells tool. The resulting spatial-data objects integrate cortical layer boundaries, Allen reference brain region annotations, PNN status, MapMyCells cell-type identities, and PNN neighborhood information. **(B)** UMAP embed-ding of the full Xenium dataset (378,349, 5 male mice, 10 sections) colored by MapMyCells class-level annotation. Cells were plotted in randomized order to avoid visual occlusion bias. Legend below. **(C)** Composition of PNN-positive (WFA+) cells by MapMyCells subclass annota-tion (confidence > 0.75). Top: the most abundant cell types among PNN+ cells are exclusively PV interneurons (red shades), with all remaining subclasses grouped as ”Other” (grey). Bottom: breakdown of the ”Other” category (n= 91) showing the top non-PV subclasses associated with PNNs. Center values indicate percentage of total PNN+ cells (top) and cell counts (bottom).

Cell type classification revealed that 97.3% of PNN-positive cells belonged to PV interneuron types across the captured areas (Figure 1B-D). The remaining 2.7% were distributed across multiple cell types with no single population enriched, suggesting technical noise rather than a biologically distinct PNN-bearing non-PV population (Figure 1C). This near-exclusive association substantially exceeds immunohistochemical estimates of 70–80% PNN on PV cells in cortex^13^. 9,265 cortical PV cells were identified, of which 34% (3,182) were enwrapped by PNNs. We next asked whether PNN-positive cells constitute a transcriptionally distinct subtype. In each sampled major brain area, Cortex, Striatum and Thalamus, we found that PV subclasses were the only consistently PNN-enwrapped subclasses (Figure 2A). In cortex, which comprises the majority of our dataset, the 052 Pvalb Gaba subclass is the dominant PNN-enwrapped subclass, whereas the PV positive chandelier cells are mostly devoid of PNNs. Subclass 052 consists primarily of PV basket cells, though it may include cells historically classified under other names (e.g. inter-laminar cells and bushy cells). We refer to this subclass as PV basket (PvB) cells throughout. In Striatum, all detected PNN+ cells belonged to the same Gabaergic Pvalb positive subclass (053 STR Prox1 Lhx6 Gaba), which was also the case for all detected PNNs in thalamus (093 RT-ZI Gnb3 GABA, Figure 2A). At the finest cell-type resolution (cluster), no single subtype dominates, although some clusters of PvB cells are enriched for PNNs (Fig 2A). UMAP embedding of neocortical PvB (L2/3 - L6a) revealed a continuous transcriptional landscape with no discrete cluster corresponding to PNN status (Figure 2B-E). Instead, both PNN presence and PNN intensity mapped as gradients across the population. This pattern was not explained by cortical layer or brain region, which showed no systematic relationship with PNN status (Figure S1). Although UMAP visualization suggested potential substructure within PNN+ neurons, density-based clustering (HDBSCAN) on the full PCA embedding found no evidence for discrete subpopulations, consistent with PNN status reflecting a continuous transcriptional axis (Figure S1E-F).

**Figure 2.**
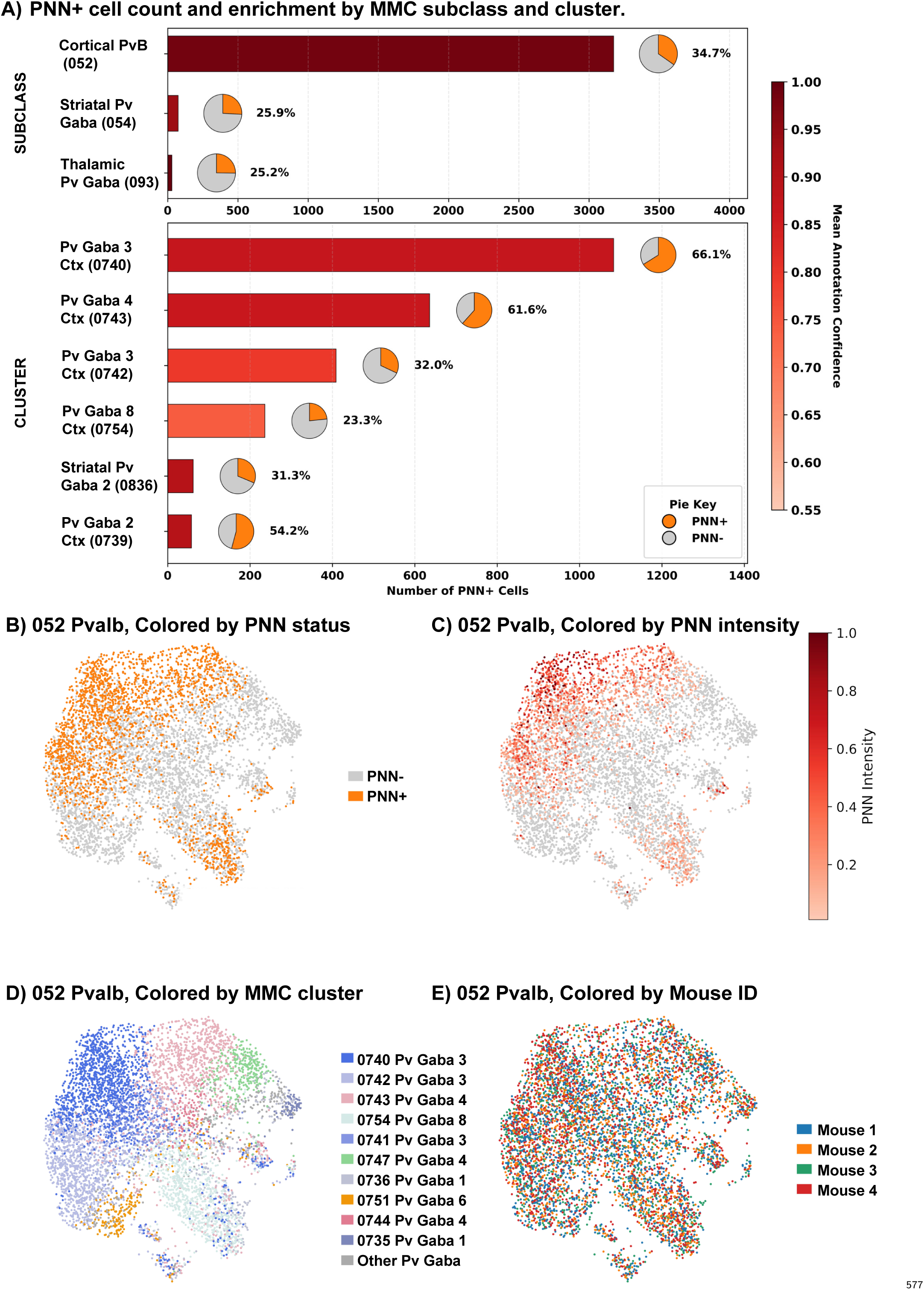
The PV basket cell subclass is strongly enriched for PNNs, but PNN+ cells do not form a distinct subcluster. **(A)** PNN+ cell counts by MapMyCells subclass and cluster colored by mean annotation con-fidence score. Inset pie charts indicate the proportion of each subclass/cluster that is PNN+ reflecting PNN enrichment. **B–E)** UMAP embeddings of the ”052 Pvalb Gaba” (PvB) subclass (n = 2260 PNN+, 6624 PNN-, 4 animals). **B)** Colored by PNN status (WFA+/WFA−), showing the distribution of PNN-bearing and PNN-lacking PvB cells in transcriptomic space. **C)** Colored by PNN fluorescence intensity (WFA signal), highlighting graded differences in PNN expression across the PvB subclass. **D)** Transcriptomic subclusters within the PV population visualized by UMAP colored by MapMyCell cluster. **E)** Colored by mouse ID, demonstrating no apparent batch effects between animals.

### Transcriptional signatures distinguish PNN-positive from PNN-negative PV neurons

To identify genes associated with PNN status within the PvB population, we performed differential expression analysis between PNN-positive and PNN-negative PvB cells. In the 297-gene Xenium panel (Table S1.1), we identified 68 differentially expressed genes (adjusted p < 0.05, |log FC| > 0.75, n=5): 22 upregulated and 46 downregulated in PNN-positive cells (Figure 3B, Table S1.2). PNN-positive neurons showed elevated expression of structural ECM components including Acan, Hapln1, and Tnr, confirming the expected molecular signature^19^, alongside canonical fast-spiking markers: Syt2, the mature NMDA receptor subunit Grin2a^21^, and Mybpc1. This signature was graded rather than binary: expression of Syt2, Adamtsl5, and Acan decreased progressively across PvB clusters with declining PNN enrichment (Figure 3A), while the least PNN-enriched cluster (0751) showed reduced Pvalb expression and elevated Calb1, Sst, and Id2, genes associated with other interneuron identities^24,25^. The most PNN-enriched cluster (0743) was distinguished by high Reln expression. Many of these genes also correlated strongly with PNN intensity within the PvB population, particularly Grin2a, but notably Pvalb expression did not (Figure 3C). Contactin-1 (Cntn1), recently implicated as a PNN cell-surface link^28^, was expressed across all Pv clusters regardless of PNN status or brain region. Of the PNN-associated ADAMTS proteases, cortical PvB cells consistently expressed Adamts8 and Adamts15 as previously reported^19^, whereas striatal PV cells with PNNs preferentially expressed Adamts5, and thalamic PV cells expressed only moderate levels of Adamts15 (Figure 3B).

**Figure 3.**
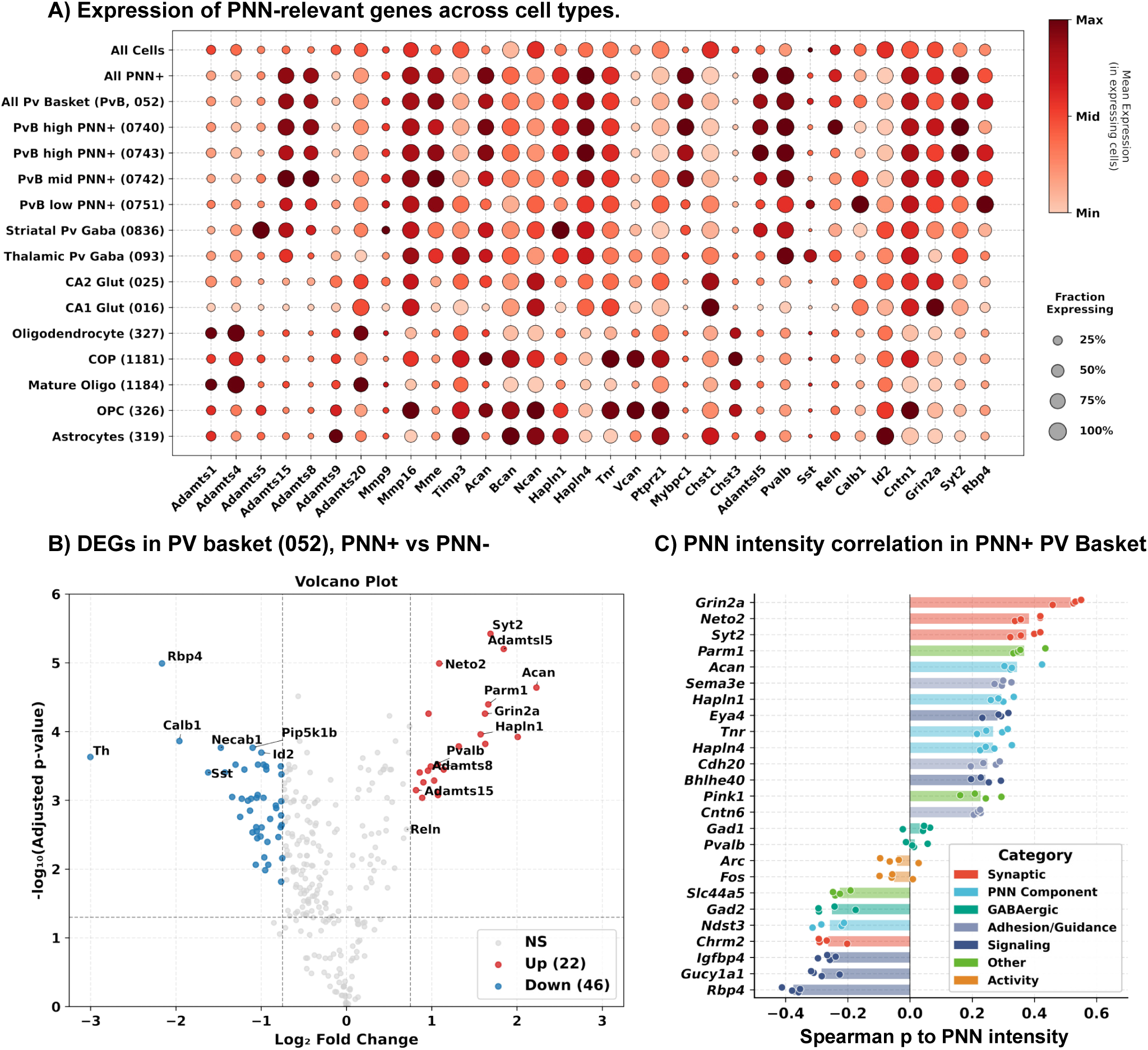
Differential gene expression across PNN+ and PNN− cells. **A)** Dotplot showing expression of PNN-associated genes across cell populations. Dot size indi-cates the fraction of cells expressing each gene; color intensity indicates mean expression level in expressing cells (scaled per gene). Gene categories include ADAMTS metalloproteinases, matrix metalloproteinases, core PNN components (Acan, Bcan, Ncan, Hapln 1 & 4, Tnr, Vcan, PtPrz1), and glutamate receptor subunits (Grin2a). **B)** Volcano plot showing log fold change versus statistical significance (−log adjusted p-value) for all 297 genes in the Xenium panel in PvB cells. Red points indicate significantly upregulated genes in PNN+ neurons; blue points indi-cate significantly downregulated genes (p-value < 0.05, |log FC| > 0.75). Differential expression was calculated using paired t-tests on pseudobulk samples aggregated per animal (n = 5 mice), with p-values adjusted for multiple testing using (Benjamini-Hochberg) Known PNN components (Acan, Hapln1) and synaptic markers (Syt2, Grin2a, Neto2) are among the most significantly upregulated genes. **C)** Top genes correlating (Spearman) to PNN intensity in PvB cells, with highlighted gene categories (n=4).

Comparison with CA2 pyramidal neurons, which bear PNN-like structures, revealed a distinct molecular profile. CA2 cells showed elevated Acan relative to CA1 but lacked the PNN+ signature of PV cells: no Adamts8 or Adamts15, elevated Timp3, and high Chst1 expression. Ncan expression was low in PV neurons but elevated in both CA1 and CA2 pyramidal cells, while Bcan was predominantly expressed in glia (COPs, OPCs, astrocytes). Glial cells, including astrocytes and oligodendrocytes, have been reported to contribute to PNN formation and maintenance^29,30^. To test whether spatial proximity to PNNs influences glial transcriptomes, we analyzed oligodendrocyte lineage cells and astrocytes in the isocortex. PNN-proximal cells showed no clustering separation from PNN-distant cells in any glial population (Figure S2). Expression of PNN-associated genes, including CSPGs, link proteins, and ECM-modifying proteases did not differ significantly between groups (paired Wilcoxon, BH-corrected; Table S1.5).

We next asked whether gene expression could predict PNN status. Using nested cross-validation with leave-one-animal-out splits, we trained a two-stage classifier using L1 regularization for feature selection and L2-regularized logistic regression to distinguish PNN+ from PNN- PV neurons^31^. The model achieved robust performance (AUC = 0.87; sensitivity = 68%; specificity = 85%; permutation p < 0.001; Figure 4A-C). Feature accumulation analysis revealed that the binary model reached 95% of plateau performance with only 4 genes, indicating PNN acquisition is predicted by a remarkably compact transcriptional signature (Figure S4 A).

**Figure 4.**
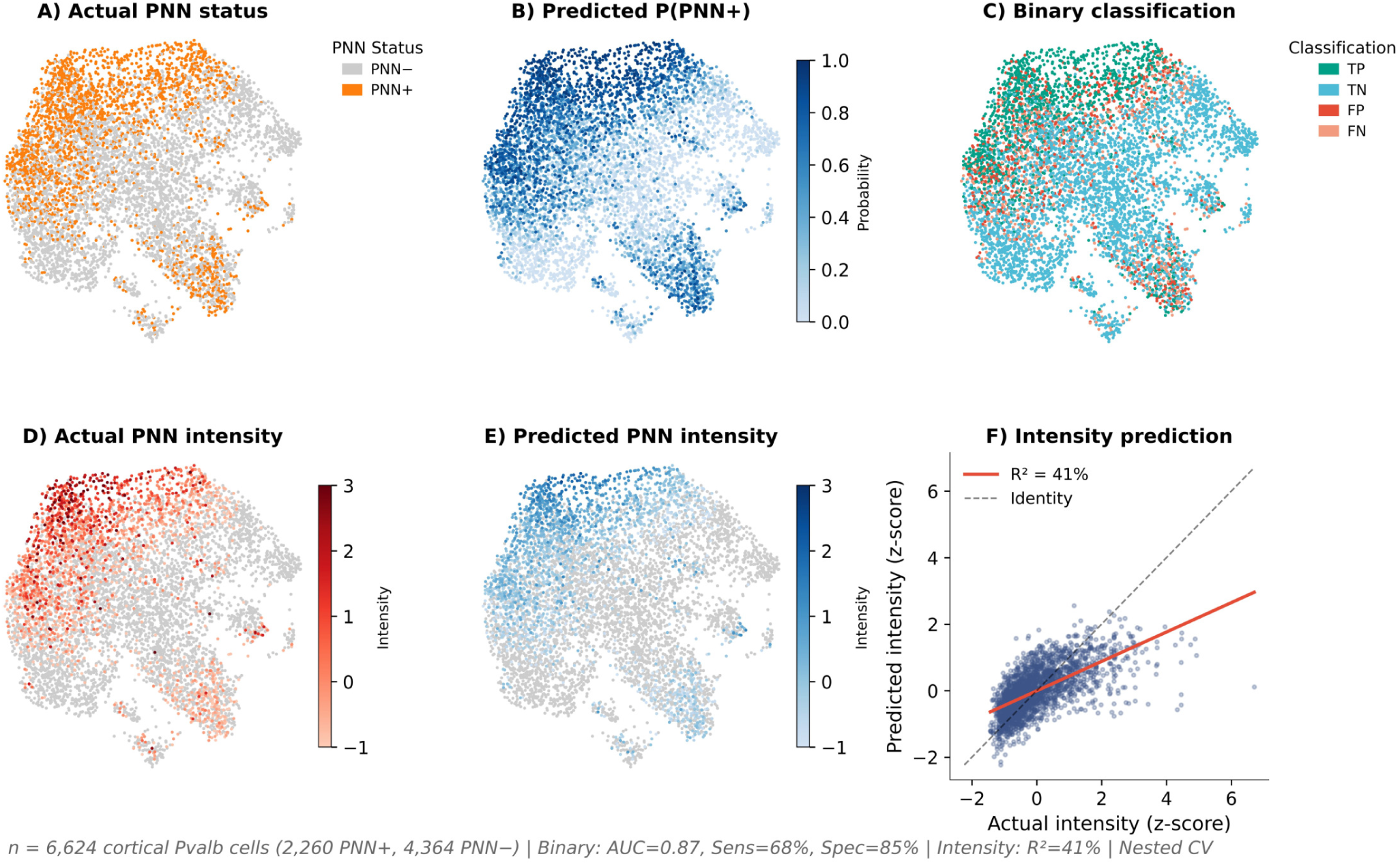
Gene expression predicts PNN status in PV basket cells. **A)** UMAP embedding of cortical PvB cells colored by PNN status. **B)** Predicted probability of PNN+ status. **C)** Binary classifier performance: true positives (TP), true negatives (TN), false positives (FP), and false negatives (FN). **D)** Same embedding colored by PNN intensity (z-scored within slice). **E)** Predicted PNN intensity (PNN+ neurons only; PNN− neurons in gray). **F)** Pre-dicted versus actual PNN intensity. Solid red line: linear regression fit; dashed line: identity. **Binary model:** LASSO feature selection followed by L2-regularized logistic regression (AUC = 0.87). **Intensity model:** LASSO feature selection followed by Ridge regression (R² = 41%). Both models evaluated using nested leave-one-animal-out cross-validation with feature selection performed independently within each training fold. n = 6,624 PV neurons (2,260 PNN+, 4,364 PNN−) from 4 mice.

The binary classifier distinguishes PNN presence from absence, but PNN intensity varies considerably among the PNN-positive cells (Figure 2). We therefore asked whether gene expression could also predict PNN intensity. Among PNN-positive cells, a regression model explained 41% of the variance in PNN intensity (R² = 0.41, permutation p < 0.001; Figure 4D-F). In contrast to the binary model, intensity prediction required 25 genes to reach 95% of plateau, suggesting the transcriptional contribution to intensity is weaker and more distributed (Figure S4 B).

Comparison of coefÏcients between the binary and intensity models revealed distinct gene categories (Table S1.4). Binary-only predictors, including Pvalb, Mybpc1, and Col19a1, strongly predicted PNN presence but not intensity, potentially marking transcriptional competence for PNN acquisition. Genes predicting both outcomes with concordant effects included Syt2, Grin2a, Hapln4, Acan, and Tnr, as well as negative predictors Gad2, Ndst3, Snap25, and Tacr1 (substance P receptor), identifying cells with lower PNN-formation probability and reduced intensity. A small set of genes (Gria4, Igfbp4, Lypd6) predicted lower PNN intensity without affecting acquisition probability. No genes showed clear opposite effects between models; none promoted acquisition while limiting intensity, or vice versa.

To test whether the cortical PNN signature generalizes across brain regions, we applied the cortex-trained classifier to hippocampal PvB cells (n = 226; 98 PNN+, 128 PNN−). The model achieved AUC = 0.77, lower than cortical performance (AUC = 0.87) but substantially above chance. This partial generalization suggests the core specialization axis is conserved, though hippocampal PvB cells may have region-specific PNN determinants not captured by the cortical signature.

### Label transfer to Allen Brain Atlas enables genome-wide differential expression analysis

While the 297-gene Xenium panel is sufÏcient to identify PNN-associated signatures and train a predictive model, it cannot reveal genome-wide transcriptomic differences. To extend our analysis, we therefore applied the trained classifier to Allen Brain scRNA-seq data, which profiles 32,285 genes across the adult mouse brain^16^.

We extracted 34,326 cortical PvB cells in the Allen dataset and applied our binary classifier to infer PNN status. For differential expression analysis, we retained 24,743 high-confidence cells (probability ≥ 0.7 for PNN+ or ≤ 0.3 for PNN−) from 69 donors with at least 10 cells per group (Figure S5.A). Differential expression using PyDESeq2 with donor and PNN status as design factors (n = 69 donors, 24,743 cells) identified 7,626 differentially expressed genes (padj < 0.001, no LFC cutoff): 2734 upregulated and 4,892 downregulated in predicted PNN-positive cells (Figure 5A, Table S1.3). Of these, 7,401 were genes absent from the Xenium panel, representing novel transcripts associated with PNN status. For downstream pathway analysis, we applied no fold-change threshold; for individual gene visualization, we highlight a |log2FC| > 0.5 threshold (Figure 5B). Effect sizes were generally modest (mean |log2FC| = 0.47, 90th percentile = 0.98), consistent with a graded transcriptional shift.

**Figure 5.**
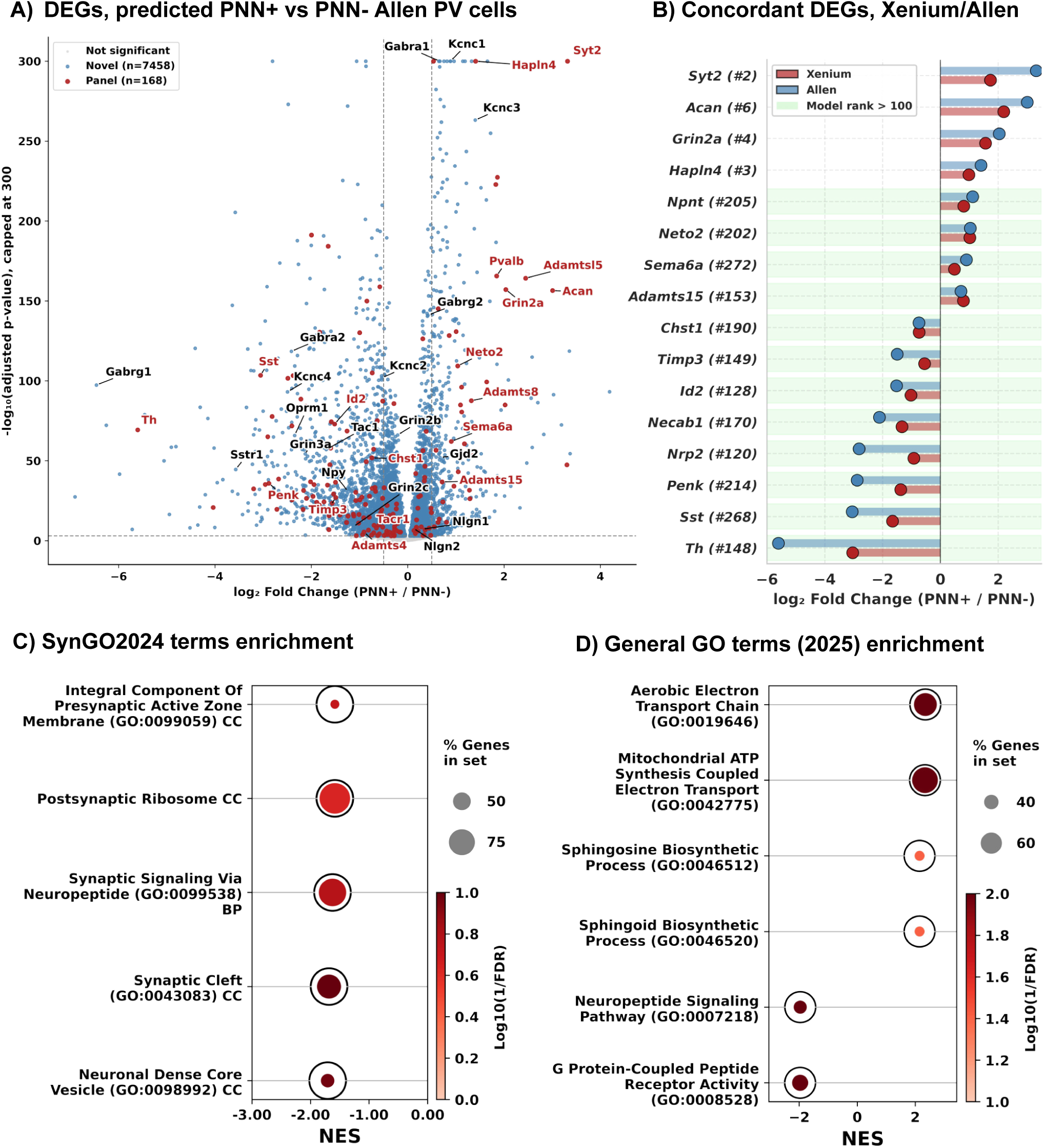
Genome-wide differential expression and pathway enrichment in predicted PNN+ versus PNN− PV neurons. **A)** Volcano plot showing differential expression between predicted PNN+ and PNN− PvB cells in Allen Brain scRNA-seq data^16^. Genes present in the Xenium panel (red) and novel genes (blue) are highlighted. Genes of interest are named. Horizontal dashed line: padj = 0.001; ver-tical dashed lines: log2FC = ±0.5. Gray points: padj ≥ 0.001. Differential expression analysis was performed using PyDESeq2 with donor and PNN status as design factors, with pseudob-ulk aggregation by donor (n = 69 donors, 24,743 cells). **B)** Lollipop plot of select genes show-ing concordant direction of effect between Xenium spatial transcriptomics and Allen scRNA-seq, spanning high and low model ranks. Xenium Log2 fold change in blue, Allen in red. Model rank > 100 highlighted in teal background. **C)** Gene set enrichment analysis using SynGO 2024 synap-tic ontology. Dot size indicates percentage of leading-edge genes; color indicates significance (−log10 FDR, inferno). NES, normalized enrichment score. **D)** Gene set enrichment analysis using general Gene Ontology sets (Molecular Function, Cellular Component, Biological Process; 2025 release). Formatting as in (C).

To validate that label transfer preserved biological signal, we examined genes with minimal influence on the classifier trained on Xenium data. Regardless of model rank, gene expression aligned well in the two datasets (Figure 5B). Among genes ranked below the top 100 by model coefÏcient, 57 were independently significant DEGs in both Xenium and Allen datasets (padj < 0.05). All of these 57 genes showed concordant direction of effect (Figure S5B). This consistency in genes that did not drive classifier training confirms that inferred PNN labels capture genuine biological differences rather than circular artifacts of the prediction model. While highly concordant, Allen data effect sizes were consistently larger than in our Xenium data (linear regression slope = 1.57, Figure S5B).

Known PNN-associated genes showed expected patterns in the Allen data: Acan (log2FC = +2.5), Hapln4 (+1.2), Syt2 (+1.8), and Grin2a (+2.0) were strongly upregulated in predicted PNN-positive cells, while Sst (-3.1) was downregulated. Among low-weight validation genes, Neto2 (model rank 202; Xenium +1.02, Allen +1.04) and Bcan (rank 283; Xenium +0.56, Allen +0.86) showed near-identical effects across platforms.

### PNN status predicts fast-spiking specialization, metabolic capacity, and neuropeptide ex-pression

Predicted PNN-positive neurons showed elevated expression of genes associated with mature, fast-spiking physiology. The NMDA receptor subunit Grin2a was strongly upregulated (log2FC = +2.04), while Grin2b, Grin2c, and Grin3a were downregulated, recapitulating the developmental Grin2a/Grin2b switch as stable adult heterogeneity^21^. GABA receptor subunit profiles diverged strikingly: PNN-positive neurons expressed Gabra1 (+0.67) and Gabrg2 (+0.42), subunits that confer faster inhibitory kinetics, while PNN-negative neurons expressed Gabra2 (-2.41), Gabrb1 (-0.93), and Gabrg1 (-6.46), subunits associated with slower decay and tonic inhibition^22^. Kv3 potassium channels, essential for rapid repolarization and high-frequency firing^20^, showed a similar pattern: Kcnc1 (Kv3.1, +0.88) and Kcnc3 (Kv3, +1.40) were upregulated in PNN-positive neurons, while Kcnc2 (Kv3.2, -0.54) and Kcnc4 (Kv3.4, -2.49) were downregulated (Figure 5A, Table S1.3). Gap junction expression also differed: Gjd2, encoding Connexin-36, was upregulated in PNN-positive neurons (log2FC = +0.67). Connexin-36 forms electrical synapses between fast-spiking interneurons and is required for gamma oscillation synchrony^23^. Extending our Xenium observation, PNN-negative neurons showed markedly elevated neuropeptide expression: Sst (log2FC = -3.05), Tac1 (-1.69, encoding substance P), and Npy (-1.23). Together, this profile positions PNN-positive neurons at the canonical fast-spiking end of the PV continuum, equipped for the precise, high-frequency inhibition that PNNs are proposed to support.

To characterize biological processes underlying the full signature, we performed gene set enrichment analysis (GSEA) against GO and SynGO databases. This confirmed the neuropeptide pattern, with ”neuropeptide signaling pathway” (FDR = 0.002) and ”neuronal dense core vesicle” (FDR = 0.11) among the most enriched terms in PNN-negative cells (Figure 5C, Table S1.7). The co-expression of Pvalb with Sst-associated neuropeptides and GABA receptor subunits suggests PNN-negative neurons may represent a stable subpopulation at the transcriptomic boundary between canonical PV and Sst interneuron identities. GSEA revealed strong enrichment of mitochondrial respiration pathways in PNN-positive cells: ”mitochondrial ATP synthesis coupled electron transport” (normalized enrichment score (NES) = +2.34, FDR = 0.004) and ”aerobic electron transport chain” (NES = +2.35, FDR = 0.009). Leading edge genes included nuclear-encoded components of complexes I, III, IV, and V (Ndufa/b, Uqcr, Cox, Atp5 families). This metabolic signature is consistent with the high energy demands of sustained fast-spiking activity characteristic of mature PV interneurons (Figure 5D). Other significant hits are included in Table S1.7 and include additional cellular respiration terms, as well as G-Protein Coupled Receptor Activity, and sphingolipid metabolism.

Finally, to test whether a binary ’master switch’ gene for PNN formation exists, we screened all 32,285 genes in the Allen Brain Atlas for binary expression patterns between predicted PNN+ and PNN- PvB cells. We defined binary expression as >80% of cells below detection threshold (count <1) in one group and <30% in the other. No gene met these criteria; the closest candidate, Mybpc1, narrowly missed with 86% of PNN- cells silent but 32% of PNN+ cells also silent (Figure S6). This confirms that PNN status reflects a graded transcriptional continuum rather than a discrete cell state.

## DISCUSSION

Previous comprehensive immunohistochemical work has reported that 20–30% of cortical PNNs surround non-PV neurons^13^, leading to the hypothesis that PNNs mark functionally diverse cell populations. Here, we combined spatial transcriptomics with PNN immunolabeling to characterize the molecular identity of PNN-bearing neurons in adult mouse cortex. Our findings challenge existing assumptions about PNN cell-type heterogeneity, reveal that PNN status reflects a continuum rather than a discrete PvB subtype, and identify PNN-negative PvB neurons as a stable subpopulation retaining molecular features of immature or plastic states.

Our transcriptomic classification found that 97% of PNN-positive cells belong to PV sub-classes, consistent with recent intersectional genetic approaches that reported 99% using Vgat-Cre/PV-Flp mice^32^. This discrepancy likely reflects the limited sensitivity of fluorescent Pvalb immunolabeling, which can miss neurons with low Pvalb protein levels. Transcriptomic cell typing integrates information across hundreds of genes, providing more robust identification across the full range of expression levels. These findings suggest the apparent heterogeneity in PNN cell-type association reported in earlier studies reflects methodological limitations rather than true biological diversity.

UMAP embedding revealed no discrete cluster corresponding to PNN status; instead, both PNN presence and intensity mapped as gradients across a continuous transcriptional manifold (Figure 2D). This has implications for how we interpret PNN function. Rather than marking a specialized cell type, PNNs appear to reflect position along a specialization axis that exists within the broader, adult PV population. PNN status is predictable from gene expression (AUC = 0.87) yet does not correspond to a discrete cluster. This suggests a threshold model: cells become competent for PNN formation upon reaching a certain transcriptional state, but that state represents a point along a continuum rather than a qualitatively distinct identity. Furthermore, the absence of binary marker genes has practical implications: simple probe panels for PNN+ cell identification are not feasible. Detection would require quantitative measurement of multiple genes and application of a classifier, rather than categorical presence/absence of a single or a few markers. Our predictive modeling supports a two-stage model of PNN regulation. The binary classifier achieved high performance with only four genes, indicating that transcriptional competence for PNN acquisition is encoded by a remarkably compact signature dominated by high levels of genes canonical to fast-spiking cells. In contrast, PNN intensity prediction required 25 genes to reach 95% of plateau predictive power and explained only 40% of the variance, suggesting intensity is modulated by additional factors. The remaining 60% of intensity variance likely reflects post-transcriptional mechanisms, including activity-dependent ECM remodeling, proteolytic processing by matrix metalloproteinases, and methodological limitations of intensity measurements. This dissociation between acquisition and intensity has implications for therapeutic strategies targeting PNNs: interventions aimed at promoting or preventing PNN formation may engage different mechanisms than those modulating the density of existing PNN structures.

PNN-positive neurons displayed a coherent receptor signature of mature, fast-spiking neurons. The Grin2a/Grin2b switch, a canonical marker of synaptic maturation during development^21^, persisted as stable adult heterogeneity, with PNN+ neurons expressing mature Grin2a and PNN- neurons retaining subunits associated with earlier stages of development. Indeed, pan-neuronal Grin2a KO results in delayed maturation of PV neurons and PNNs^33,34^. Kv3 potassium channels showed a similar pattern: Kcnc1 and Kcnc3, which enable the fast repolarization required for high-frequency firing^20,35^, were elevated in PNN+ neurons, while Kcnc2 and Kcnc4 were reduced. Gap junction expression also diverged: Gjd2, encoding Connexin-36, was upregulated in PNN+ neurons. Connexin-36 mediates electrical coupling between fast-spiking interneurons and is required for gamma oscillation synchrony^23^. This combined profile suggests PNN+ neurons are tuned for the precise, temporally sharp inhibition characteristic of canonical fast-spiking PvB cells, while PNN- neurons, with their slower GABA receptor subunits and reduced Kv3.1/3.3 expression, may provide broader, less temporally precise inhibition. PNN- PV cells, which also express more Zcchc12 and 6330403K07Rik, genes associated with slow spiking, may preferentially innervate dendrites in contrast to the canonical axo-somatic PvB cells^18^.

Consistent with earlier observations that PNN-enwrapped cortical PV neurons lack Sst co-expression^19^, predicted PNN-negative PV neurons showed markedly elevated neuropeptide expression. These cells expressed 8-fold higher somatostatin, 3-fold higher substance P (Tac1), and 2-fold higher NPY compared to predicted PNN+ neurons. This was accompanied by GABA receptor subunit profiles characteristic of Sst rather than PV interneurons: elevated Gabra2, Gabrb1, and Gabrg1 versus the Gabra1/Gabrg2 signature of PNN+ cells. Transcriptomic studies have established that Sst interneurons co-express Gabra5, Gabra2, and Gabrb1, while PV interneurons co-express Gabra1, Gabrb2, and Gabrg2^22^. Our data show this same divergence within the PV population, stratified by PNN status.

The co-expression of PV with Sst-associated neuropeptides and receptors suggests PNN-neurons occupy a transcriptomic position at the boundary between canonical PV and Sst interneuron identities. This parallels continuous transcriptional variation identified in CA1 interneurons, where a single continuous variable predicted expression of fast-spiking machinery, metabolism, and GABA synthesis genes versus neuropeptides and slower receptor kinetics, correlating with axon target location from soma to distal dendrites^18^. In primary visual cortex (V1), position along the same transcriptomic continuum across all interneuron types predicted state-dependent activity: cells at the fast-spiking end of a continuum were preferentially active during synchronized, stationary brain states, while cells at the opposite end fired more during locomotion and arousal^36^. If PNN status maps onto this axis, as our data suggest, PNN+ and PNN- PV neurons may be differentially recruited depending on behavioral state. The substantial overlap with our PNN-associated signatures suggests these functional and structural axes may be linked.

Whether PNN- PV neurons represent a distinct lineage or a plastic subpopulation capable of transitioning to the PNN+ state remains to be determined. Rather than reflecting a single maturation endpoint, PNN heterogeneity may partition PV interneurons into functionally complementary populations: one providing the temporally precise inhibition required for gamma synchrony, the other maintaining flexibility for circuit adaptation. The elevated oxidative phosphorylation machinery of PNN-positive neurons reinforces this distinction. Fast-spiking interneurons are among the most metabolically demanding cells in the brain, firing at sustained high frequencies that require constant ATP regeneration^37^. The coordinated upregulation of nuclear-encoded mitochondrial genes (complexes I, III, IV, and V) suggests PNN+ neurons have adapted their metabolic infrastructure to support this demand. Conversely, PNN- neurons showed elevated ribosomal protein expression, potentially reflecting greater ongoing protein synthesis capacity consistent with a more plastic or adaptable state. Future functional characterization of PNN- PV neurons could leverage intersectional viral strategies, for example, Sst promoter-driven Cre-dependent constructs in PV-Cre mice, or vice versa, to selectively target this population without the developmental lineage confounds inherent to constitutive intersectional genetics, ie. PV-cre X Sst-Flp.

CA2 pyramidal neurons are a well-known exception to the near-exclusive association between PNNs and inhibitory interneurons^38^. In our Xenium data, CA2 pyramidal cells showed elevated Acan relative to CA1 but lacked the transcriptional signature that stratifies PV interneurons by PNN status: no Adamts8 or Adamts15, and distinct lectican composition (Ncan rather than Acan-dominant). Grin2a expression is high in glutamatergic CA2 neurons, but this is also the case for glutamatergic neurons in CA1. This suggests that while Acan expression is required for PNN assembly, the upstream transcriptional programs are cell-type specific, likely reflecting distinct functional demands for PNN formation across neuronal classes. The ADAMTS proteases implicated in PNN remodeling showed regional specificity: Adamts15 was expressed in PNN-bearing PV neurons across cortex, thalamus, and striatum, while Adamts8 was restricted to cortex. Stri-atal PV neurons instead expressed Adamts5, suggesting region-specific mechanisms of PNN turnover.

Glial cells, including astrocytes and oligodendrocytes, express several PNN-associated genes and have previously been proposed to contribute to PNN formation^39,40^. However, the extent to which glial-derived components incorporate into the mature perisomatic PNN structure remains unclear. Aggrecan, the core structural CSPG of WFA-positive PNNs, appears to be expressed predominantly by neurons^41^, and PNNs are abolished when Acan is selectively knocked out in PV cells^10^ or pan-neuronally^42^. Our spatial transcriptomic analysis aligns with these observations: glial cells near PNNs showed no differential expression of PNN components or modulators compared to distant glia, suggesting that glial cells do not adopt a specialized transcriptional program related to local PNN maintenance. These findings point to neurons as the primary source of their own PNN.

Whether specific neuronal activity drives PNN formation, and if so which patterns, remains unknown. While PNN maturation correlates with the development of gamma oscillations, it remains unclear whether gamma-frequency firing itself is required, as opposed to general neuronal activity levels or participation in network oscillations. This question has not yet been tested at single-cell resolution. Single-cell targeted optogenetics combined with longitudinal PNN imaging could address whether stimulating individual PNN-negative PV neurons at specific frequencies is sufÏcient to induce PNN formation, and whether this shifts cells along the transcriptional axis described here.

Together, these findings suggest that PNN status in the adult brain marks a molecular boundary between two stable PV configurations: one defined by fast receptor and ion-channel profiles, elevated metabolic capacity, and circuit-stabilizing ECM enwrapment, and another retaining neuropeptide signaling, developmental receptor subunits, and greater biosynthetic capacity. Whether these states arise from divergent developmental trajectories, ongoing adult plasticity, or stable circuit specialization remains unresolved. Resolving this will be critical for therapeutic approaches that aim to modulate cortical plasticity through PNN manipulation, as the answer determines whether PNN-negative PV neurons represent a targetable reservoir of circuit flexibility or a fundamentally distinct neuronal identity.

### Limitations of the study

Several limitations should be considered when interpreting these findings. First, our label transfer approach inherently propagates uncertainty: classifier predictions on Allen data reflect Xenium-defined boundaries, and cells with intermediate probabilities (which we excluded) may represent biologically meaningful states. The high-confidence thresholds (≥0.7 or ≤0.3) are conservative but discard approximately half of PV neurons. That said, the 98% directional concordance among low-weight genes supports genuine biological signal rather than circular artifacts.

Second, the receptor subunit differences we report (Gabra1/2, Grin2a/b, Kcnc1/3) predict distinct electrophysiological properties, but we did not directly measure synaptic kinetics, firing patterns, or gamma entrainment in PNN+ versus PNN- neurons. Similarly, the elevated oxidative phosphorylation signature in PNN+ cells awaits confirmation by metabolic measurements.

Third, as in most of the literature, we define PNNs by WFA reactivity. WFA labeling is sensitive to post-translational modification patterns and may not cover the full PNN population - some of which would be labeled by Acan immunostaining, but not WFA^43,44^. Nevertheless, this is a source of false negatives rather than false positives, meaning our classifier may underestimate the PNN+ population but should not spuriously inflate it.

Finally, our analysis is restricted to neocortex (mainly) in male mice, and our Xenium data was collected from a single age-group (7 months). PNN distribution and PV heterogeneity vary across brain regions and may differ by sex, or age. Age of the mice in the Allen dataset was P53 to P71, substantially younger, but still past the PNN development window. The Allen dataset includes both sexes, which partially mitigates this concern for the transferred analysis but does not substitute for direct spatial validation in female tissue. Applying the classifier to datasets spanning additional ages and both sexes would test whether the PNN-associated transcriptional axis is stable across these variables.

## METHODS

### Animals and tissue preparation

All work with experimental animals was performed at the animal facility at the Department of Biosciences, Oslo, Norway, in agreement with guidelines for work with laboratory animals described by the European union (directive 2010/63/EU) and the Norwegian Animal Welfare Act from 2010. The experiments were approved by the National Animal Research Authority of Norway (Mattilsynet, FOTS #26727). The animals were housed in groups of four to five prior to experiments in GM500 IVC cages, with ad libitum access to food and water. For enrichment, the cages had a running wheel, nesting material, and chew blocks made from wood.

Adult C57BL/6 mice (P220, n = 5 male) were used for all experiments. Mice were briefly anesthetized with isoflurane and decapitated. Dissected brains were cut along the midline and fresh-frozen in dry-ice chilled iso-pentane in Tissue-Tek® O.C.T. Compound (Sakura), in custom vacuum-formed cryo-molds (REF github) and stored at -80°C. Sagittal sections (10 µm thickness) were cut on an NX-70 cryostat and mounted on Superfrost slides for quality control, and Xenium slides for Xenium. Full sagittal sections were collected intermittently as the brain was cut lateral to medial, these sections were later used for reference atlas alignment. At 1°.5 mm ML, the OCT-embedded brain was scored with a razor blade on the chuck, to eliminate cerebellum, midbrain and the olfactory bulb, as well as excess OCT. Sections for Xenium were collected at 1°.5 to 1°.7mm ML. Using paintbrushes, tissue was separated from the frozen OCT to minimize the footprint on the xenium slides, before being cold-mounted on the xenium slides. Xenium slides were contained within the cryostat at -20 until sections from all five animals had been collected. Finaly, the xenium slides were shipped on dry ice to the Single Cell Genomics Core at the University of Eastern Finland for Xenium spatial transcriptomics.

### Spatial transcriptomics

Xenium In Situ (10x Genomics) was performed using a custom 297-gene panel comprising the standard 10x Genomics Mouse Brain Panel (247 genes) plus 50 custom probes targeting PNN-associated genes (Table S1.1). Probes were hybridized to target RNA, followed by ligation and enzymatic amplification to generate multiple copies of each RNA target, as described in the Xenium Probe Hybridization, Ligation, Amplification, and Cell Segmentation Staining User Guide (User Guide CG000749). The prepared Xenium slides were then loaded onto the Xenium Analyzer for imaging and analysis, following the Xenium Analyzer User Guide (User Guide CG000584). Instrument software version 3.2.1.2 and analysis software version 3.2.0.7 were used.

### WFA immunofluorescence

Following Xenium imaging, sections were stained for perineuronal nets using biotinylated Wisteria floribunda agglutinin (WFA). Sections were blocked in 3% BSA, 0.1% Triton-x, incubated with biotinylated WFA (1:200 of stock, overnight at 4C, Vectorlabs), washed, and incubated with streptavidin-conjugated Alexa 647 (1:500 of stock, overnight 4C, ThermoFisher). Parvalbumin and Acan (SWANT PV-27 and Milipore ab1031) immunostaining was attempted but did not yield any signal, likely due to the specific tissue processing required for Xenium (10um, fresh frozen) or issues stemming from Xenium chemistry. Sections were mounted in SlowFade Glass (ThermoFisher) and imaged on a spinning-disc confocal microscope (20× objective, Olympus SpinSR, Yokogawa CSU-W1 confocal spinning disk). WFA images were integrated as an image layer in the spatial data objects following alignment using Xenium Explorer and Sopa^45^.

### Image registration

Slice images were aligned to the Allen Mouse Brain Atlas (CCFv3) using the QUINT workflow (QuickNII/VisuAlign)^27^. Briefly, 1°0 full saggital sections spaced 2°00 um apart were collected on superfrost slides prior to the collection of saggital hemi-sections for Xenium. These full brain slices, along with the final Xenium slices, were used to register to the allen reference atlas. Registered reference atlas label images were aligned to the Xenium data in Napari instances of the SpatialData objects. Registration of label images enabled assignment of anatomical region labels to individual cells in the Xenium dataset.

### Cell segmentation and data integration

Default Xenium cell segmentation was refined using ProSeg^46^ with original Xenium boundaries as priors. Parameters were adjusted to minimize merged cells (–max-transcript-nucleus-distance reduced from default 60, to 10). Each tissue section was represented as a SpatialData object containing transcript coordinates, cell boundaries, and aligned WFA images (10 sections from 5 animals in total).

### PNN annotation and quantification

PNN-positive cells were identified by manual annotation in a napari instance of the SpatialData object^47^. The annotator was blind to transcriptomic results and could only see the ProSeg cell boundaries and the aligned (SOPA^45^/Xenium Explorer) WFA staining image. PNN+ cells lacking a ProSeg boundary were excluded as they are not registered cells in the Xenium dataset. Point labels marking PNN centers defined by the annotator were integrated automatically in each object (slice). A cell was classified as PNN-positive if its ProSeg-segmented boundary contained a PNN point label. For PNN-positive cells, intensity was quantified by extracting a square region (120 × 120 pixels) centered on the ProSeg boundary centroid. Mean WFA intensity was normalized to local background measured in a surrounding square (10th percentile, 360 × 360 pixels).

### Cell type classification

Cell types were assigned using the Allen Brain Atlas MapMyCells pipeline^26^ with raw transcript counts as input. Cells were classified hierarchically at multiple taxonomic levels (class, subclass, supertype, cluster). AnnData^48^ objects from all sections were merged into a single dataset. Quality filters removed cells with less than 75 transcripts, or less than 45 genes detected, as well as transcripts with QV below 20. The final dataset comprised 378,349 cells. Data were normalized (counts per 10,000) and log1p-transformed for downstream analysis.

### Differential expression analysis

*Xenium dataset.* Differential expression between neocortical (L2/3-L6a) PNN-positive and PNN-negative PV neurons was performed using paired t-tests on pseudobulked samples (n = 4 animals; one animal excluded due to poor WFA staining intensity). DESeq2 was not used as reliable dispersion estimation requires n ≥ 6 samples per group^49^. For each gene, mean log-normalized expression (log1p of CP10K-normalized counts) was calculated per animal separately for PNN+ and PNN− cells, and paired t-tests assessed the consistency of differential expression across animals. Log fold changes were computed from means back-transformed to linear scale. P-values were adjusted using Benjamini-Hochberg correction. Genes were considered differentially expressed at adjusted p < 0.05 and |log FC| > 0.75.

*Allen Brain dataset.* To enable genome-wide differential expression analysis, we applied the trained binary classifier to 34,326 PV basket cells (subclass 052) in an Allen Brain scRNA-seq data^16^, and inferred PNN status using classifier probabilities. For classification, raw counts were log -transformed and z-scored. High-confidence cells were retained using probability thresholds of ≥0.7 (PNN+) or ≤0.3 (PNN−). Donors with fewer than 10 cells in either group were excluded, yielding 24,743 cells from 69 donors. Raw counts were summed per donor per condition to create pseudobulk samples. Differential expression was performed using PyDESeq2^50^ with donor and PNN status as design factors. Genes with fewer than 10 counts in fewer than 10 samples were excluded. Log fold changes were shrunk using the apeGLM method^51^. Genes were considered differentially expressed at Benjamini-Hochberg adjusted p < 0.001.

*Gene set enrichment analysis.* Gene set enrichment analysis (GSEA) was performed using gseapy prerank^52^ on genes ranked by Log fold change, with 1,000 permutations. Gene sets were obtained from SynGO^53^ (2024 release) and Gene Ontology (GO Biological Process, Molecular Function, and Cellular Component; 2025 release via Enrichr^54^). Minimum and maximum gene set sizes were 5 and 1,000, respectively. Terms were considered significant at FDR < 0.25^55^.

### Machine learning models

*Data preprocessing.* Gene expression and PNN intensity values were z-scored within each tissue section to control for technical variation across slices.

#### Binary classification (PNN presence)

To predict PNN status from gene expression, we used a nested cross-validation framework to prevent data leakage. The outer loop used leave-one-animal-out cross-validation (n = 4 folds). Within each outer fold, feature selection was performed on training data only using L1-regularized logistic regression (LogisticRegressionCV, scikit-learn^31^; 5-fold inner CV for regularization parameter selection). The top 30 genes by absolute coefÏcient were retained, and an L2-regularized logistic regression model was trained on these features. Predictions were generated for held-out test data. Final performance (AUC = 0.87, sensitivity = 68%, specificity = 85%) was calculated from concatenated out-of-fold predictions. Statistical significance was assessed by permutation testing (n = 1,000 permutations, p < 0.001). Of the top 30 genes per fold, 21 were selected in all 4 folds.

#### Intensity regression (PNN structure density)

Among PNN-positive cells, we applied the same nested cross-validation framework using LassoCV for feature selection and Ridge regression on the top 30 selected features. Model performance (R² = 0.41) was calculated from out-of-fold predictions and assessed by permutation testing (p < 0.001). Gene selection was stable (19/30 genes in all folds), the top-ranked genes were consistently selected across all folds.

#### Feature accumulation analysis

To assess signature compactness, genes were ranked by abso-lute coefÏcient magnitude from L1-regularized models fit on the full dataset. Models were then trained using the top k genes (k = 1 to 200) with performance evaluated by leave-one-animal-out CV. The binary model reached 95% of plateau performance with 4 genes; the intensity model required 25 genes.

#### Label transfer to Allen Brain Atlas

The trained binary classifier was applied to Allen Brain Atlas scRNA-seq PV basket cells (n = 34,326) to infer PNN status. Input features were limited to genes present in both datasets. Predicted probabilities were thresholded at 0.5 for classification; high-confidence subsets (probability ≥0.7 or ≤0.3) were used for differential expression analysis.

## RESOURCE AVAILABILITY

### Lead contact

Requests for further information and resources should be directed to and will be fulfilled by the lead contact, Sverre Grødem.

### Materials availability

This study did not generate new materials.

### Data and code availability

All original code will be deposited at Github (github.com/Sverreg/PNNcontinuum2026) upon pub-lication and is publicly available as of the date of publication. All original data will be made avail-able through Ebrains. Any additional information required to reanalyze the data reported in this paper is available from the lead contact upon request.

## ACKNOWLEDGMENTS

This work was funded by University of Oslo, Research Council of Norway Grant No.352863 to M.F, the Norwegian Health Association, and Research Council of Norway Grant No.325892 to T.H. We acknowledge the Single Cell Genomics Core at the University of Eastern Finland, supported by Biocenter Kuopio and Biocenter Finland, for providing spatial transcriptomics services. Finally, the authors thank all members of the lab for their support.

## AUTHOR CONTRIBUTIONS

*Conceptualization and design, S.G and G.H.V; methodology, S.G; investigation, S.G; writing-–original draft, S:G; writing-–review & editing, all authors; funding acquisition, M.F, T.H; resources, M.F*

## DECLARATION OF GENERATIVE AI AND AI-ASSISTED TECH-NOLOGIES

*During the preparation of this work, the authors used Claude Opus 4.5 and Gemini 2.5 Pro to assist with data processing, analysis, visualization and writing. After using this tool or service, the authors reviewed and edited the content as needed and take full responsibility for the content of the publication*.

---

**Figure S1.**
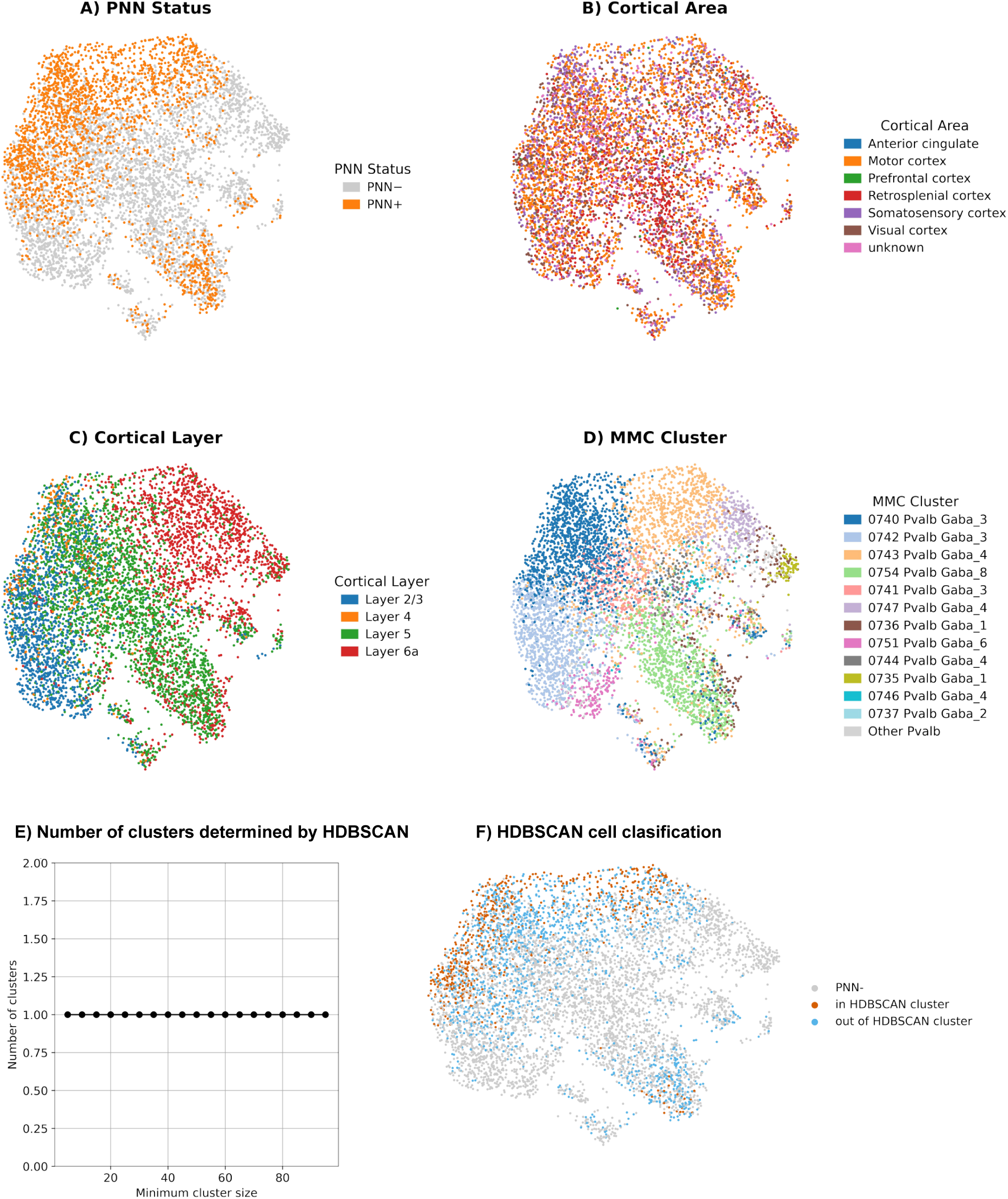
**Spatial and transcriptomic distribution of PNN+ PV interneurons.** UMAP projections of PV neurons (n = 6,624 cells from 4 mice) colored by **A)** PNN status, **B)** cortical area, **C)** cortical layer, and **D)** MapMyCells cluster assignment (top 12 clusters by cell count shown; remaining clusters grouped as ”Other Pvalb” in grey). PNN+ neurons do not segregate by cortical area or layer, suggesting that PNN association is driven by cell-intrinsic molecular identity rather than anatomical location. **E)** The number of clusters determined through HDBSCAN^56^ on PCA coordinates of all PNN+ cells across different values of the minimum cluster size parameter. A single cluster was consistently identified. **F)** UMAP projection colored by the clustering label of PNN+ cells with HDBSCAN (min_cluster_size = 10).

**Figure S2.**
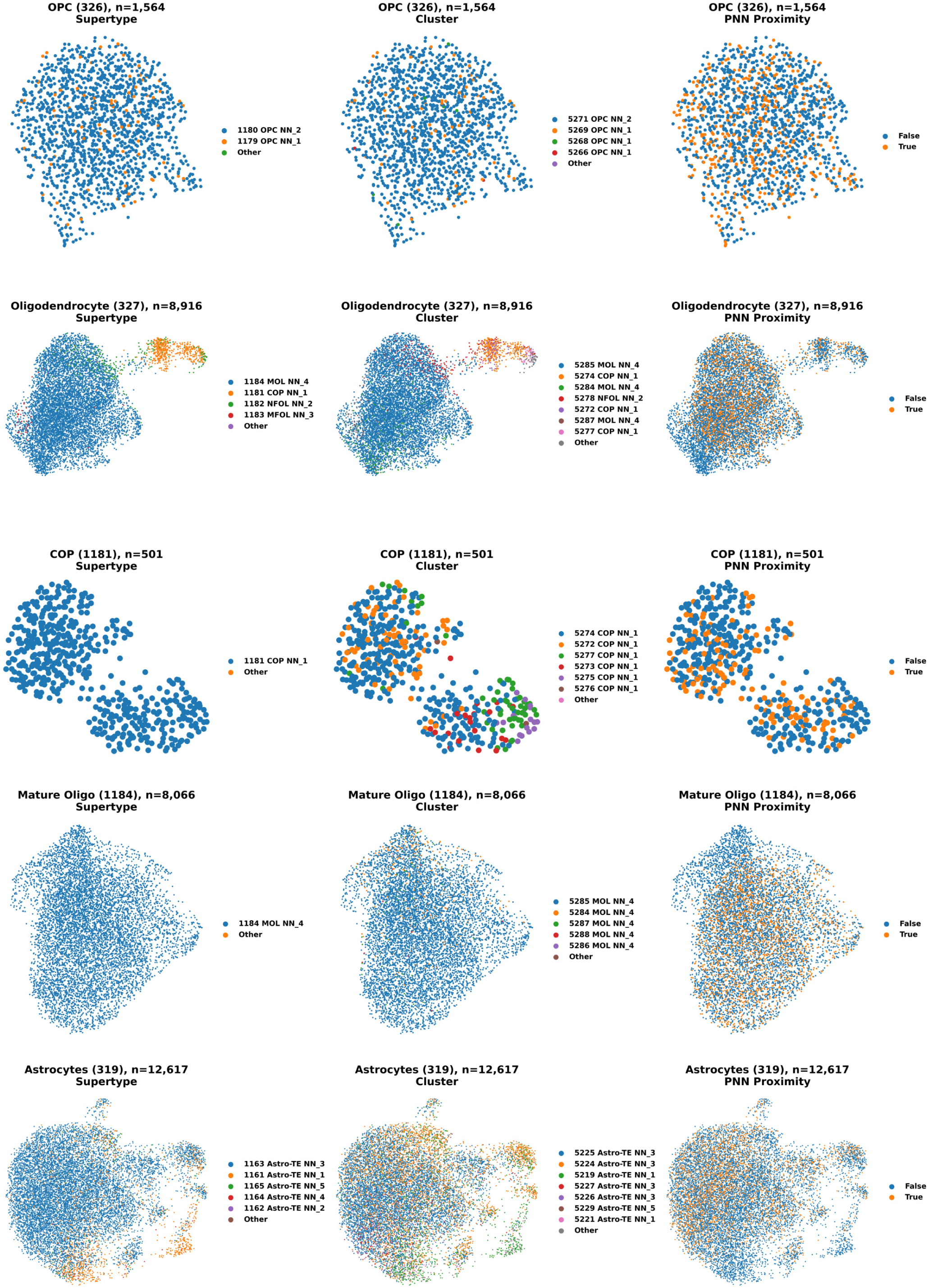
UMAP visualization of glial cell types in relation to PNN proximity. UMAP projections of isocortical glial populations (Layers 2–6b) computed independently for each cell type using the top 200 highly variable genes (Xenium). Rows show OPCs (326), oligodendrocytes (327), committed oligodendrocyte precursors (COP, 1181), mature oligodendrocytes (MOL, 1184), and astrocytes (319). Columns display coloring by transcriptomic supertype (left), cluster (middle), and PNN proximity status (right). PNN-proximal cells were defined as cells within 50 μm of a WFA-positive cell, identified using squidpy’s spatial neighbors graph (radius = 50 μm); Cell counts per group are indicated in panel titles.

**Figure S3.**
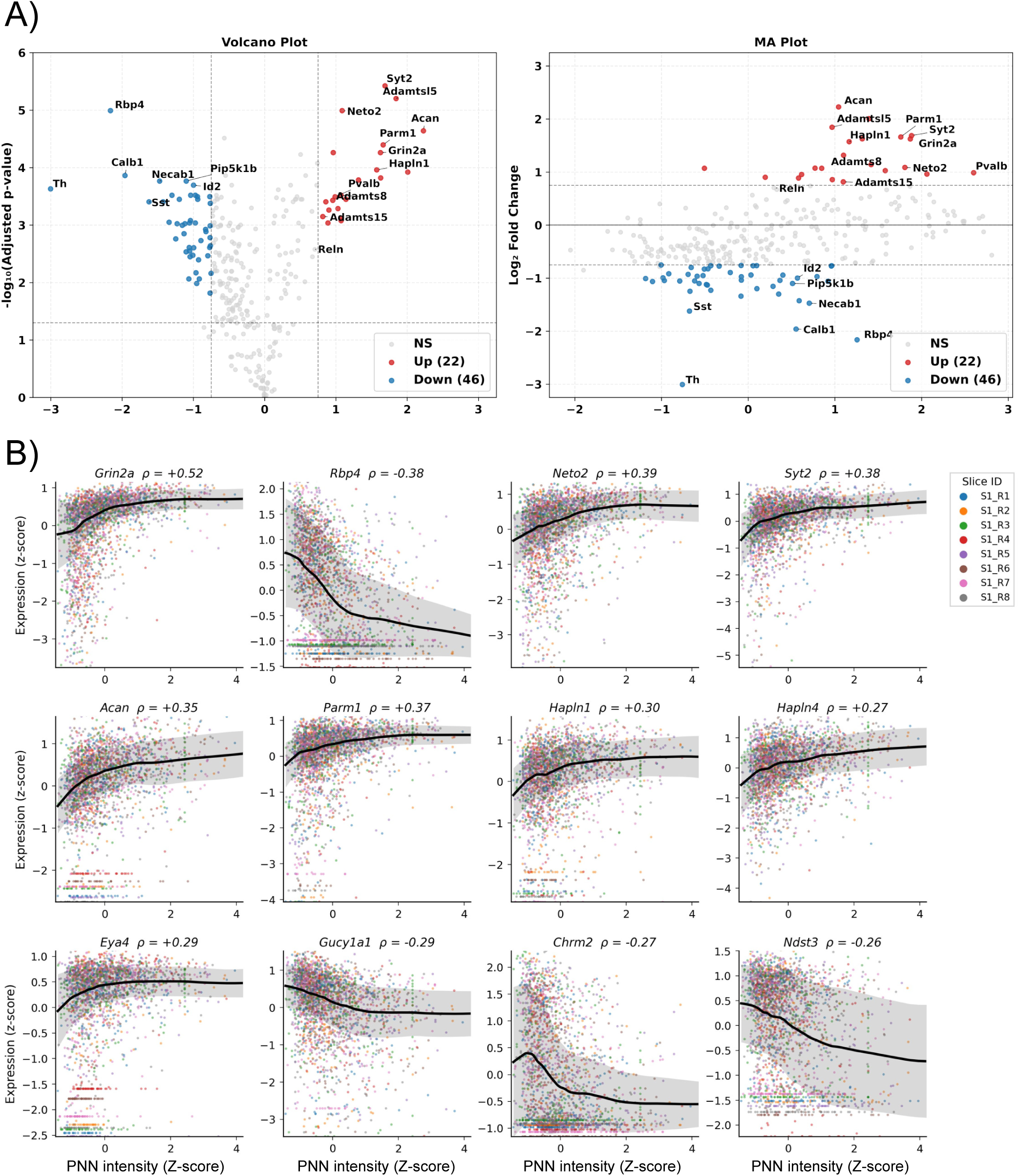
MA plot of differential gene expression between PNN+ and PNN− PV interneurons and scatterplots with Lowess smoothed trend of gene expression vs PNN intensity. **A)** MA plot showing log fold change versus mean expression (log) for all 297 genes in the Xenium panel. Red points indicate significantly upregulated genes in PNN+ neurons; blue points indicate significantly downregulated genes (p-value < 0.05, |log FC| > 0.75). Differential expression was calculated using paired t-tests on pseudobulk samples aggregated per animal (n = 4 mice), with p-values adjusted for multiple testing using the Benjamini-Hochberg method. Known PNN components (Acan, Hapln4) and synaptic markers (Syt2, Grin2a, Neto2) are among the most significantly upregulated genes. **B)** Scatterplots of expression vs PNN intensity for top genes correlating with PNN status. Both gene expression and PNN intensity were z-scored within each slice to control for batch effects. Points are colored by slice (n=8 slices from 4 mice), demonstrating consistency across biological replicates. Black line shows LOWESS smoothed trend (frac=0.3); shaded region indicates ±1 SD. Spearman ρ shown for each gene. Analysis restricted to cortical layers 2/3–6a (n = 2,345 PNN+ PvB cells).

**Figure S4.**
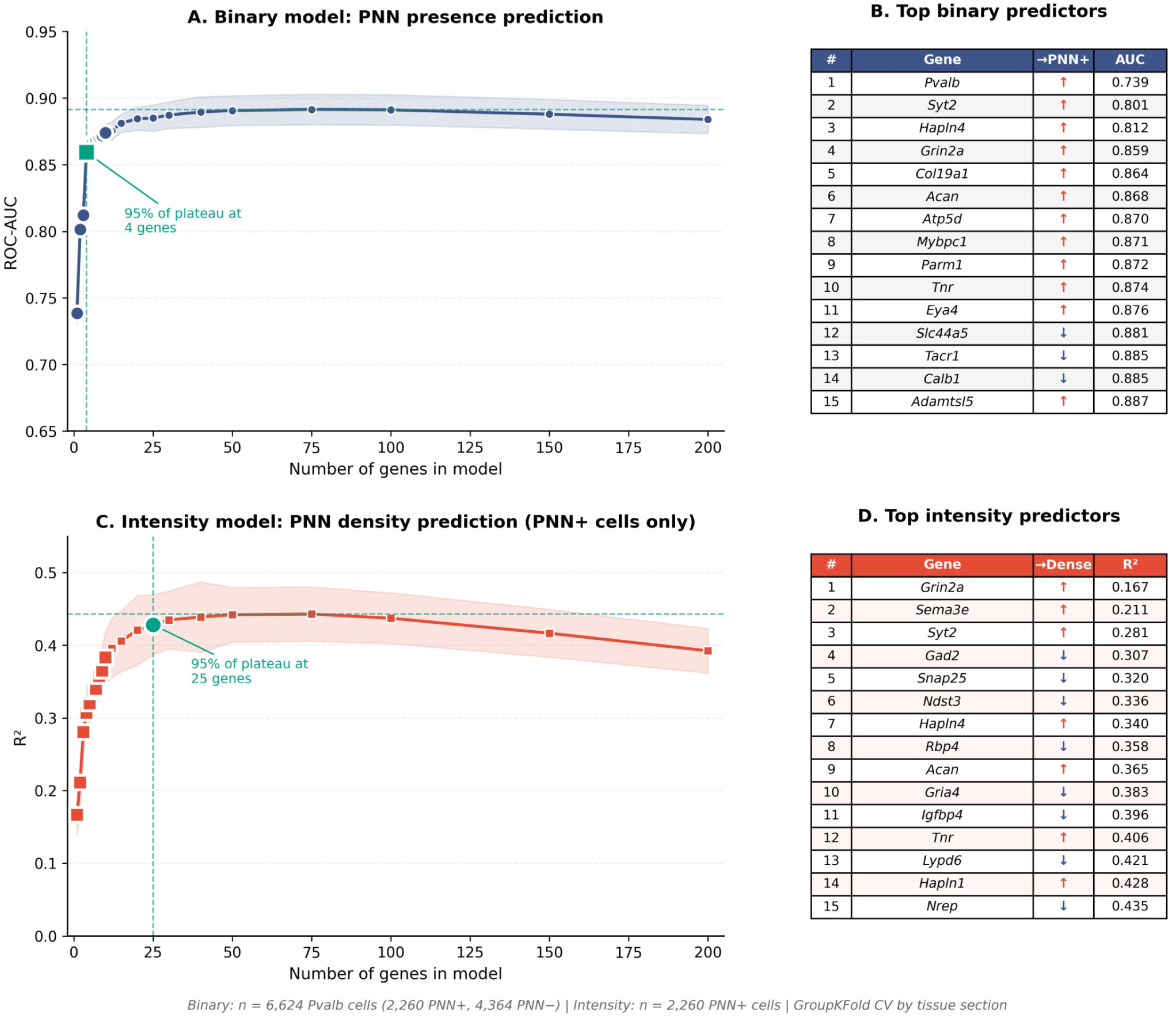
Feature accumulation analysis reveals minimal gene sets sufÏ-cient for PNN status prediction. **A)** Classification performance (ROC-AUC) as a function of the number of genes included in the binary model predicting PNN presence. Genes were ranked by absolute LASSO coefÏcient and added incrementally. Performance evaluated using 4-fold cross-validation grouped by animal. Shaded region indicates ±1 SD. Teal marker indicates 95% of maximum performance (4 genes). **B)** Top 15 binary model predictors ranked by absolute coefÏcient weight. **C)** Regression performance (R²) for the intensity model predicting PNN density among PNN+ neurons. Performance plateaus at 2°5 genes (R² = 0.42). **D)** Top 15 intensity model predictors. **Binary model**: n = 6,624 PV neurons (2,260 PNN+, 4,364 PNN−). **Intensity model**: n = 2,26 PNN+ neurons. Cross-validation grouped by animal ID (n = 4 mice).

**Figure S5.**
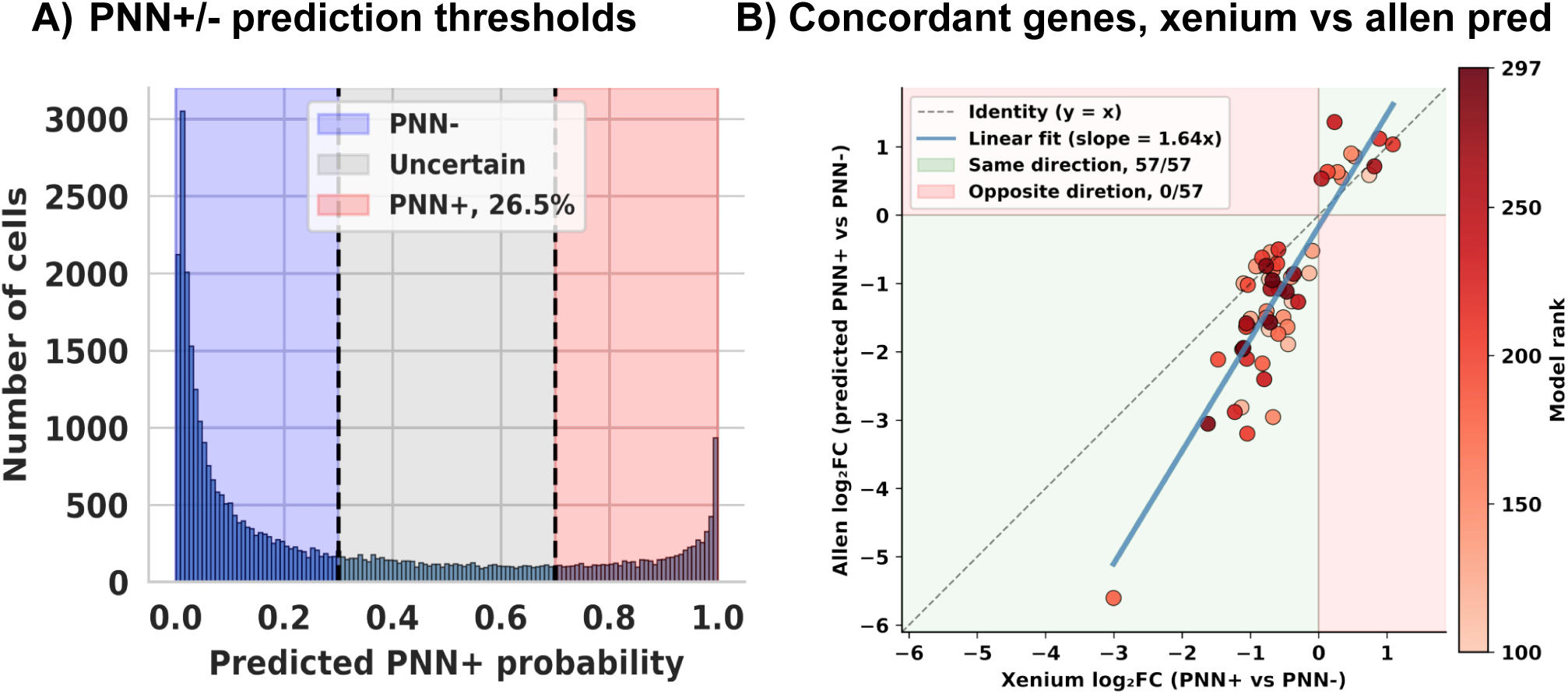
Label transfer validation and cross-platform concordance. **A)** Distribution of predicted PNN+ probability for Allen Brain PvB cells (n = 34,326). Dashed vertical lines indicate thresholds for high-confidence classification: cells with probability ≥0.7 were classified as PNN+, cells with probability ≤0.3 as PNN−. Cells with intermediate probabilities (gray region) were excluded from differential expression analysis. **B)** Concordance of differential expression between PNN+/PNN- in Xenium spatial transcriptomics and predicted PNN+ vs PNN-in Allen scRNA-seq for genes with low classifier influence (model rank > 100). Each point represents a gene significant in both datasets (padj < 0.05); color indicates classifier rank (darker = lower rank, less influence on predictions). Green shaded quadrants indicate concordant direction of effect; red shaded quadrants indicate discordance. Of 57 genes significant in both datasets, 57 showed concordant direction. Solid blue line: linear regression fit (slope = 1.64); dashed line: identity (slope = 1).

**Figure S6:**
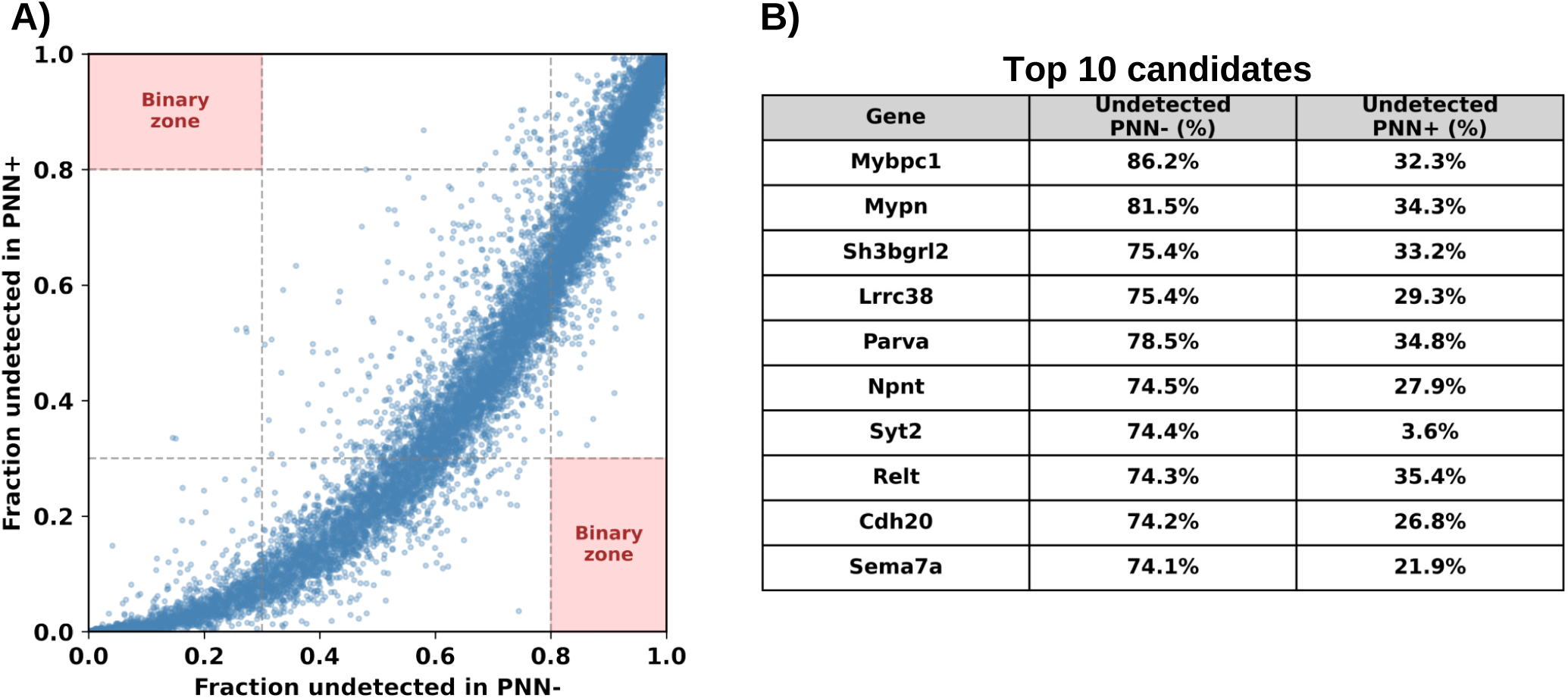
No binary marker gene distinguishes PNN+ from PNN− PV neu-rons. **B)** Scatter plot of fraction undetected in PNN+ (y-axis) versus PNN− (x-axis) neurons for all 32,285 genes in Allen Brain Atlas scRNA-seq data. Red shaded regions indicate binary expression criteria: >80% undetected in one group and <30% in the other. No gene falls within these regions. **C)** Top 10 candidate genes ranked by proximity to binary criteria. Mybpc1 approached but did not meet the threshold (86% undetected in PNN−, 32% undetected in PNN+). Data from predicted PNN+ and PNN− PV neurons (probability threshold ≥0.7 or ≤0.3, n = 24,743 cells total).

## SUPPLEMENTAL TABLES INDEX

All tables below are consecutive sheets in the attached Table_S1.xlsx file.

**Table S1.1.** Xenium gene panel.

Table S1.2. Differential gene expression between PNN+ and PNN- PvB cells in Xenium dataset.

**Table S1.3.** Differential gene expression between predicted PNN+ and PNN− PvB cells in Allen Brain scRNA-seq data.

**Table S1.4.** Ranked binary model coefÏcients.

**Table S1.5.** Differential gene expression in glial cells by PNN proximity.

**Table S1.6.** Distribution of cells per brain area in Xenium dataset.

**Table S1.7.** Gene ontology analysis results.

